# Diagnostic Use of PCR in Carbapenamase-producing Enterobacteriaceae (CPE): An Improved Method to Overcome the presence of inhibitors for DNA Extraction from Blood Cultures

**DOI:** 10.1101/312686

**Authors:** Zeti Norfidiyati Salmuna, Murnihayati Hassan

## Abstract

**Background:** Carpanenamase-producing *Enterobacteriaceae* (CPE) has emerged as a threat to hospitalized patients. Phenotypic test such as Modified hodge test was less sensitive and specific especially to detect *bla*NDM-1 which is the most predominant genotype in this region. Nucleic acid amplification technology offers improved specificity and sensitivity. However, there is limitation in this method which is failed amplification due to the presence of inhibitors. In this study, we try to use previous method described by Villumseen et al with some modification using another DNA extraction kit.

**Methodology/Principle findings:** Three confirmed isolates of *bla*NDM-1 carbapenamase-producing *Klebsiella pneumoniae* was spiked with 10 mls sterile whole blood in an aerobic Bactec Plus until the growth was detected. The blood specimen was taken and was subjected to DNA extraction method using two commercialized kits followed with multiplex PCR. All the three isolates revealed *bla*NDM-1 genotypes. Internal control was amplified in all three isolates.

**Conclusion/significance:** This study proved that we can get a specific and early diagnosis of CPE by using nucleic acid amplification technique directly from blood culture bottle and eliminate the effect of inhibitors.

## Introduction

The increasing prevalence of extended spectrum beta-lactamases (ESBL) *enterobacteriaceae*, made carbapenem a preferred drug for treatment that results in the emergence of carbapenem-resistant *Enterobacteriaceae* (CRE) that has become a formidable threat to public health(28). Beside limiting treatment option, infections due to CRE are associated with higher morbidity and mortality rates (3, 23). Higher mortality rates due to carbapenemase-producing *enterobacteriacea* (CPE) are multifactorial such as underlying diseases, lack of effective antimicrobials therapy and delay in starting effective antimicrobials (18).

The prevalence of CRE in South East Asia was about 2.8% as documented in one study (13). However, a study in Uganda showed that carbapenamase prevalence was 22.4% by phenotypic test, but slightly higher with the genotypic test which was 28.6% (16). However, there are limited studies regarding incidence of CRE genotype in Malaysia. One study done among 13 isolates of carbapenem-resistant *Klebsiella pneumniae* (CRKP) in a Malaysia university hospital found that only two, *bla*NDM1 and *bla*IMP4 genotypes were detected (11).

There are mainly two different mechanisms that are responsible for carbapenem resistance in *Enterobacteriaceae* which is either ESBL or AmpC enzymes hyper production combined with porin loss or alteration or upregulated efflux pump and or production of carbapenem-hydrolyzing carbapenamases (6, 7, 20, 22, 26).

Carbapenamases are categorized into Ambler molecular classification as follows, the Class A carbapenamases (GES, KPCs) which are inhibited by clavulanic acid; the Class B or metallo-β-lactamases (VIM, IMP, NDMs) which are inhibited by ethylene diamine tetra-acetic acid (EDTA); and the Class D oxacillinases which are not affected by clavulanic acid or EDTA (8, 10, 19, 21, 27).

The most important and common cause of carbapenem resistance in CRE is due to plasmid-mediated carbapenamase production (1, 12, 21, 22, 27) that have resulted in multiple outbreaks reported from various regions of the world.

CRE is defined as Enterobacteriaceae that are resistant to any carbapenem antimicrobial (i.e:minimum inhibitory concentration of ≥4 mcg/ml for doripenem, meropenem or imipenem OR ≥2 mcg/ml for ertapenem) **OR** documented to produce carbapenamase (by MHT, Carba-NP or genotype) (4)

The Clinical and Laboratory Standard Institute (CLSI) recommended modified Hodge test (MHT), Carba-NP and genotypic test for confirmation of carbapenamase production (5). Rapid detection of this resistance mechanism is important for prescription of appropriate antimicrobial therapy and infection control measures. Although genotypic test (molecular method) is the gold standard (2), but due to budget constraint, it is only available in few reference laboratories.

Multiplex real-time PCR has proven its superiority over conventional culture and phenotypic method. A multi-centre evaluation of real-time multiplex PCR for detection of carbapenamase genes showed that the sensitivity and specificity reached 100% and it is a robust, reliable and rapid method for detection of the most prevalent carbapenamases (24). Another study showed that the overall sensitivity and specificity of multiplex PCR assay was 99% and 100% respectively (17).

Limited studies showed that direct PCR can shorten time of detection of CRE compared to culture (14). Direct PCR proved useful in outbreak settings and high prevalence areas where rapid carbapenamase detection is critical especially for infection control management (15).

In a hospital, blood cultures (BCs) are usually collected from septic patients and send before the antibiotic is given. BCs act as a close system, therefore contamination is very unlikely. However, the blood culture bottles contain inhibitors of the PCR that require additional steps to overcome the inhibitors. Sensitivity of the assay to yield the CRE genotypes is highly dependent on the DNA-recovery.

Sodiumpolyanetholesulfonate (SPS) that serves as an anti-coagulant in blood culture bottle has become an inhibitor of the PCR as it tends to copurify with DNA (9). Therefore, there are several studies done to prevent PCR inhibition. Our study is very unique as we are able to use the previous methods and apply it with Nucleospin extraction kit. Surprisingly the PCR was not inhibited and we are able to yield the CRE genotypes implicated in the isolates tested.

## Materials and Methods

This method was done to 3 confirmed CRE isolates (*bla*NDM-1 carbapenamase-producing *Klebsiella pneumoniae*) labelled as A1, A2 and A3 from previous study. The colonies were then spiked on blood culture bottles (BACTEC^TM^ aerobic Plus) using 10 mls sterile whole blood and incubated in BACTEC machine. After the blood culture bottle gave the ‘beep’ sound, 100 μl of the positive blood culture specimen was then used for DNA extraction methods.

### DNA extraction methods

Method 5 (M5) DNA extraction methods that were used in previous study were used in this study (25).

#### Method 1 (M1) : using DNeasy^®^ Blood & Tissue Kit (Qiagen, Hilden, Germany)

##### a. Separation phase

A 100 μl aliquot of the positive blood culture specimen was mixed with 100 μl lysis buffer [5 M UltraPure^TM^ guanidine hydrochloride] (Invitrogen, CA) in 100 mM UltraPure^TM^ Tris-Hcl (Ph 8.0; Invitrogen, Paisley, UK) and 10 μl proteinase K (20 mg/ml; Qiagen) and incubated 10 minutes at room temperature. A total of 600 ml of ultrapure water (Invitrogen) was added followed by 800 ml 99% benzyl alcohol (Reagent Plus H, Sigma-Aldrich, Brondy, Denmark) and mixed. In order to separate the phases, the sample was then centrifuged at 20,000 x g for 5 minutes at room temperature.

##### b. DNA extraction phase

The 200 μl supernatant (the aqueous phase) was then transferred to a new tube. No proteinase K was added and the sample was not incubated during the lysis step. The final elution step was done in 100 μl buffer AE. This method is suitable for use with all blood culture media.

#### Method 2 (M2): using Nucleospin^®^ Blood QuickPure (Macherey-Nachel, Düren, Germany)

##### a. Separation phase

The method for separation phase was the same as in Method 1.

##### b. DNA extraction phase

The 200 μl supernatant (the aqueous phase) was then transferred to a new tube. No proteinase K was added and the sample was not incubated during the lysis step. The final elution step was done in 50 μl buffer AE.

### Experiment : Multiplex PCR

Multiplex real-time PCR amplification for the simultaneous detection of NDM-1, VIM, IMP, OXA-48 and KPC genes are done using commercially available LightMix Modular IMP, KPC, OXA-48, VIM and NDM-1 Carbapenamase multiplex kit ( by Tib Molbiol, Germany; licensed for local distribution under Roche) has been carried out on a CFX96 Touch Bio-Rad Real-Time PCR machine following the kit protocol.

Each carbapenamase gene will be detected according to target wavelength as described in the kit by the manufacturer.

#### Ethical considerations

Ethical approval was obtained from the USM Research and Ethics Committee before this study was commenced (Reference number: USM/JEPeM/ 16120537).

## Results

The results for this experiment was shown in Table 1 below.

**Table 1.** The effectiveness of DNA extraction methods, M1 and M2 in recovering *bla*NDM-1 carbapenamase-producing *Klebsiella pneumoniae*

| **DNA-extraction method** | **Positive for *bla*NDM-1** | **Internal control (IC) amplification** | **Non-template control (NTC) amplification** |
| --- | --- | --- | --- |
| **M1** |  |  |  |
| A1 | + | + | - |
| A2 | + | + | - |
| A3 | + | + | - |
| **M2** |  |  |  |
| A1 | + | + | - |
| A2 | + | + | - |
| A3 | + | + | - |
| M1: Qiagen, DNeasy® Blood & Tissue Kit |  |  |  |
| M2: Macherey-Nachel, Nucleospin® Blood QuickPure |  |  |  |

Table 1 showed the effectiveness of DNA extraction methods, M1 and M2 in recovering *bla*NDM-1 carbapenamase-producing *Klebsiella pneumoniae.* Both kits demonstrated amplification of internal control and multiplex PCR revealed *bla*NDM-1 carbapenamase-producing *Klebsiella pneumoniae*.

## Discussion

This study has its unique feature as we are able to use the method previously described (25) and mixed it with the Method 2 by using Nucleospin^®^ Blood QuickPure for DNA extraction from blood culture bottle. The results obtained are comparable for the three isolates. These findings showed 100% sensitivity and specificity. The first person that effectively solve the problems of inhibitors from blood cultures (BCs) were Fredrick and Relman in which they identified anti-coagulant sodium polyanetholesulfonate (SPS) presence in BCs (9). We found out that by adding Proteinase K treatment in lysis step, increasing the centrifugation speed to 20000 x g as well as adding extra water in the separation phase has solved the carry-over of inhibitors to the final DNA preparation in M2.

Currently, the diagnosis of CPE relies on phenotypic method (modified-hodge test) and Carba NP. But with these findings, it suggests that most cases of severe CPE can be diagnosed early by using PCR.

